# SPLiCR-seq: A CRISPR-Based Screening Platform for RNA splicing Identifies Novel Regulators of IRE1-XBP1 Signaling Under ER Stress

**DOI:** 10.1101/2025.05.20.655206

**Authors:** Qianqian Ying, Yongchen Chen, Luochen Shen, Yang Xu, Ruilin Tian

## Abstract

RNA splicing is fundamental to cellular function, yet systematic investigation of its complex regulation has been limited by existing methods. Here, we present SPLiCR-seq (<u>SPL</u>icing regulator identification through <u>CR</u>ISPR screening), a high-throughput CRISPR screening platform that enables direct measurement of RNA splicing outcomes for pooled genetic perturbations, overcoming limitations of traditional fluorescence-based approaches. Applying SPLiCR-seq to investigate *XBP1* splicing during the unfolded protein response (UPR), we conducted targeted and genome-wide screens across diverse cellular contexts, revealing both common and cell-type specific regulators. Notably, we identified GADD34 (*PPP1R15A*) as a novel modulator of IRE1-XBP1 signaling, demonstrating that it directly interacts with IRE1 and functions independently of its canonical role in eIF2α dephosphorylation. Pharmacological inhibition of GADD34 using Sephin1 effectively suppressed *XBP1* splicing and alleviated CAR-T cell exhaustion in an *ex vivo* model, leading to enhanced tumor-killing capacity across multiple cancer models. This work not only establishes a powerful new tool for systematically studying RNA splicing regulation but also uncovers a promising therapeutic strategy for improving CAR-T cell immunotherapy through modulation of the IRE1-XBP1 pathway.

## Introduction

RNA splicing is an essential process in eukaryotic cells that ensures the precise removal of non-coding introns from precursor mRNA transcripts and the joining of protein-coding exons to produce functional mature RNA[1]. This process enables the generation of diverse protein isoforms from a single gene and contributes to complex cellular functions and responses. Dysregulation in RNA splicing is linked to numerous diseases, including cancer, neurological disorders, and immune dysfunction[2–5].

Conventional RNA splicing occurs in the nucleus and is mediated by the spliceosome, a highly dynamic complex of proteins and small nuclear RNAs that recognize specific splicing signals. The spliceosome precisely modulates splicing outcomes based on cell type and environmental cues, ensuring proper gene regulation. In addition to canonical nuclear splicing, RNA molecules can also undergo unconventional splicing in response to specific cellular stress signals. A notable example is the unconventional splicing of X-box binding protein 1 (XBP1) mRNA, which occurs in the cytoplasm during the unfolded protein response (UPR)[6]. The UPR is triggered when misfolded or unfolded proteins accumulate in the endoplasmic reticulum (ER), creating stress on the cell’s protein-folding machinery[7,8]. Under these stress conditions, the transmembrane protein IRE1α (inositol-requiring enzyme 1 alpha, encoded by the ERN1 gene), a key UPR sensor, becomes activated and catalyzes the cytoplasmic excision of a 26-nucleotide segment from unspliced *XBP1* (*XBP1u*) mRNA[6,9]. This excision induces a frameshift, producing the mature, spliced *XBP1* (*XBP1s*) mRNA, which encodes the transcriptionally active XBP1 protein. This protein functions as a potent transcription factor, upregulating the expression of genes involved in protein folding, ER-associated protein degradation (ERAD), and lipid biosynthesis, thereby enabling cells to restore ER homeostasis[10]. In cancer, the IRE1-XBP1 axis emerges as a critical regulator of tumor progression and immune evasion[11]. In tumor cells, it supports adaptive survival under stress and metabolic reprogramming, creating an immunosuppressive microenvironment[12–14], while in immune cells, it contributes to T cell dysfunction and exhaustion[15–17]. Therapeutic strategies targeting this axis hold significant potential for improving cancer immunotherapy outcomes.

Recent advances in CRISPR-based genetic perturbation technologies have enabled large-scale screens to identify regulators of specific splicing events. However, existing genetic screening approaches for RNA splicing often rely on fluorescent or luminescent reporter systems[18–30], where splicing outcomes are converted into fluorescent or luminescent protein expression. These methods have several limitations. First, they provide an indirect measurement of splicing, which lack precision and temporal resolution, as the fluorescent signal lags behind actual RNA splicing events. Second, they rely on fluorescence-activated cell sorting (FACS), restricting application in primary cells, organoids, or other contexts where sorting is impractical. These challenges hinder the study of dynamic and context-specific RNA splicing regulation.

To overcome these limitations, we developed SPLiCR-seq (<u>SPL</u>icing regulator identification through <u>CR</u>ISPR screening), a CRISPR-based screening platform that directly assesses RNA splicing phenotypes by next-generation sequencing (NGS) for pooled genetic perturbations. By eliminating the need for fluorescent reporters and FACS, SPLiCR-seq provides a more direct, efficient, and precise approach for identifying splicing regulators across diverse biological settings. The simplicity of SPLiCR-seq enabled us to conduct multiple large-scale screens to investigate *XBP1* splicing during UPR across various cellular contexts, including diverse cell types, ER stressors, and treatment timepoints. These screens revealed numerous known and novel regulators of *XBP1* splicing under different conditions. Among our key findings, we discovered that GADD34 directly interacts with IRE1α to modulate *XBP1* splicing under ER stress. Furthermore, we demonstrated that pharmacological inhibition of GADD34 reduces CAR-T cell exhaustion and enhances their tumor-killing capacity, highlighting the therapeutic potential of targeting GADD34 in cancer immunotherapy.

## Results

### Development and Validation of the SPLiCR-seq Vector

The SPLiCR-seq platform was designed to link sgRNA-induced genetic perturbations with direct RNA splicing readouts via NGS. To achieve this, we engineered the SPLiCR-seq vector based on a previously published CROP-seq vector (pMK1334)[31,32]. The SPLiCR-seq vector includes a U6-driven sgRNA expression cassette for CRISPR-based gene perturbation and a splicing reporter designed to monitor the splicing level of a specific event of interest. The splicing reporter is co-transcribed with the U6-sgRNA sequence under the control of an EF1α promoter. Using paired-end NGS, the sgRNA identity is captured from one read and the splicing phenotype of the reporter is captured from the other, enabling precise mapping of genetic perturbations to their impact on RNA splicing (Fig. 1A).

**Fig. 1:**
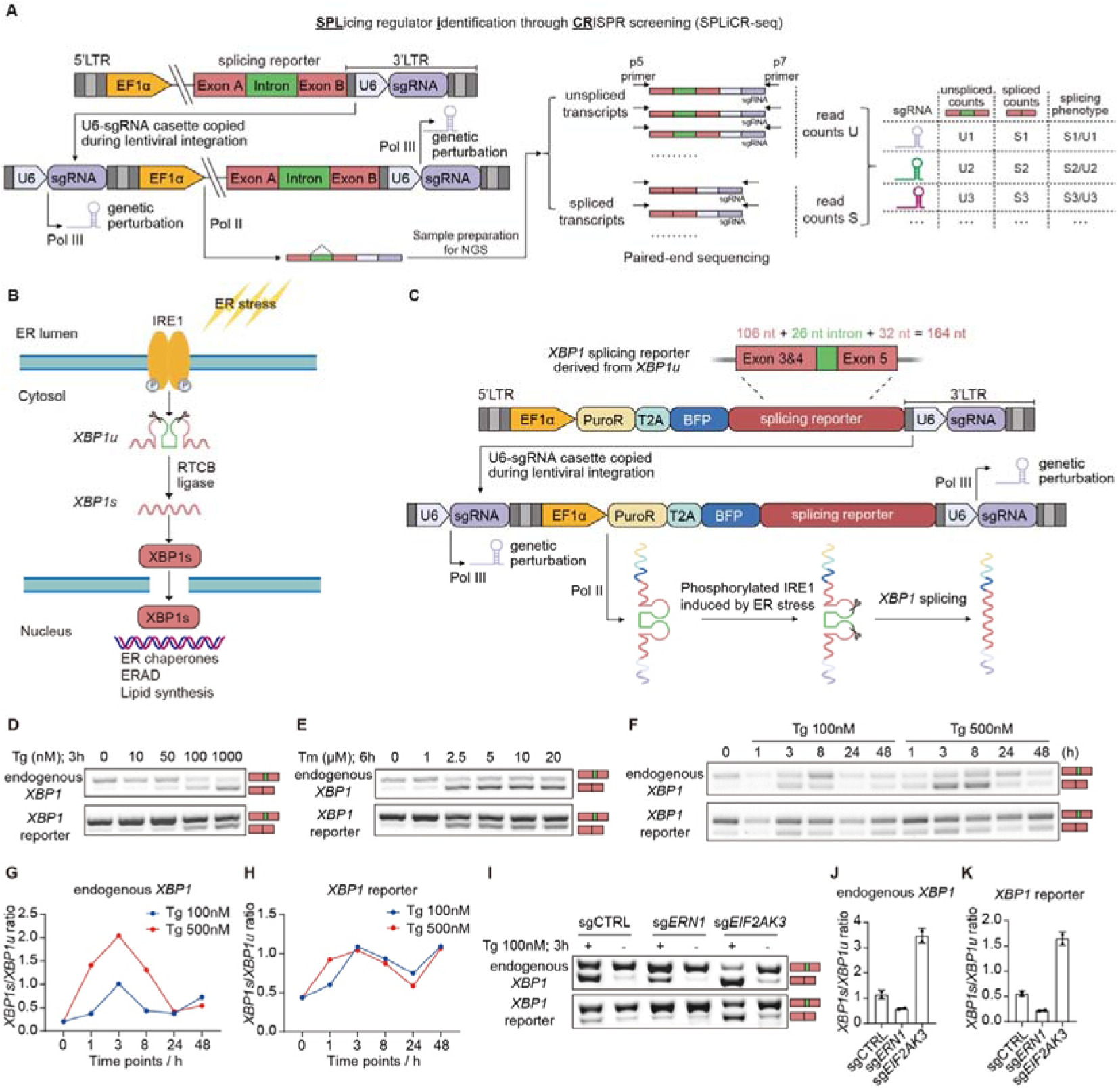
Development and Validation of the SPLiCR-seq Vector. **(A)** Schematic of the SPLiCR-seq platform. The SPLiCR-seq vector enables the co-transcription of a splicing reporter and sgRNA driven by the EF1α promoter, while sgRNA expression for gene perturbation is driven by the U6 promoter. Paired-end sequencing links the splicing phenotype of the reporter to the corresponding sgRNA identity. **(B)** Diagram of the IRE1/XBP1 pathway. Under ER stress, IRE1’s RNase domain cleaves unspliced *XBP1* mRNA (*XBP1u*), removing a 26-nucleotide intron. The resulting spliced *XBP1* mRNA (*XBP1s*) is translated into a potent transcription factor that induces the expression of ER quality control genes to mitigate stress. **(C)** Design of the SPLiCR-seq vector for *XBP1* splicing. The reporter contains the 26-nucleotide intron and 106-nucleotide 5’ and 32-nucleotide 3’ flanking sequences from *XBP1u*. **(D, E)** Dose-dependent splicing of endogenous *XBP1* and the *XBP1* reporter in the SPLiCR-seq vector in HEK293T cells treated with increasing concentrations of either Tg for 3 hours (**D**) or Tm for 6 hours (**E**), as assessed by RT-PCR. Spliced and unspliced products are indicated. **(I) (F)** Time-course analysis of *XBP1* splicing dynamics for endogenous *XBP1* and the *XBP1* reporter in the SPLiCR-seq vector in HEK293T cells treated with Tg (100 nM or 500 nM) for the indicated durations, as assessed by RT-PCR. Spliced and unspliced products are indicated. **(G, H)** Quantification of time-course splicing levels for endogenous *XBP1* (**G**) and *XBP1* reporter (**H**) using grayscale analysis of RT-PCR bands. **(I)** Representative RT-PCR analysis showing splicing of endogenous *XBP1* and *XBP1* reporter in SPLiCR-seq vectors expressing a control, *ERN1* or *EIF2AK3* sgRNA in CRISPRi-HEK293T cells. **(J, K)** Quantification of endogenous *XBP1* splicing (**J**) and *XBP1* reporter splicing (**K**) from (**I**) (mean ± SD, n = 2 biological replicates).

To demonstrate the utility of SPLiCR-seq, we applied this platform to study the unconventional splicing of *XBP1* during UPR (Fig. 1B). For this purpose, we incorporated an *XBP1* splicing reporter into the SPLiCR-seq vector. This reporter contains the 26-nucleotide intron and flanking sequences from unspliced *XBP1* (*XBP1u*) mRNA, which undergo IRE1-mediated cleavage under ER stress conditions (Fig. 1B, C).

We first validated whether the *XBP1* splicing reporter in the SPLiCR-seq vector could accurately capture the splicing dynamics of endogenous XBP1 under ER stress. Cells expressing the reporter were treated with increasing concentrations of the ER stress inducers thapsigargin (Tg) or tunicamycin (Tm)[33–35]. We found that the splicing of the reporter closely mirrored the dose-dependent splicing of endogenous *XBP1* mRNA, with both showing increased splicing at higher concentrations of Tg (Fig. 1D) and Tm (Fig. 1E). Similarly, time-course experiments demonstrated that the reporter splicing followed the same temporal dynamics as endogenous *XBP1*, with splicing levels increasing with treatment time from 0 to 8 hours and subsequently declining by 24 hours, consistent with previously reported *XBP1* splicing patterns[9] (Fig. 1F-H, Fig. S1A, B).

Next, we evaluated whether the SPLiCR-seq vector could reliably detect the effects of genetic perturbations on *XBP1* splicing. To this end, we cloned sgRNAs targeting two well-characterized regulators of the IRE1-XBP1 pathway into the SPLiCR-seq vector: *ERN1*, which encodes the enzyme IRE1 responsible for *XBP1* splicing, and *EIF2AK3*, which encodes PERK, a kinase that mediates a parallel UPR pathway whose inhibition is known to enhance IRE1-XBP1 signaling[36,37]. The vectors were transduced into a HEK293T cell line (CRISPRi-HEK293T) engineered to stably express the CRISPR interference machinery (dCas9-BFP-KRAB) for gene knockdown. As expected, *ERN1* knockdown significantly reduced the splicing of both endogenous *XBP1* and the reporter, while *EIF2AK3* knockdown showed the opposite effect (Fig. 1I-K). These results demonstrate that the SPLiCR-seq vector accurately reflects endogenous splicing dynamics and can reliably monitor the effects of genetic perturbations on splicing.

### RBP-focused SPLiCR-seq Screens Identify Cell Type-Specific Regulators of *XBP1* Splicing

Next, we performed large-scale SPLiCR-seq screens to identify regulators of *XBP1* splicing in two distinct cellular contexts: HEK293T cells, representing transformed cells, and induced pluripotent stem cells (iPSCs), representing normal cells with a normal karyotype. We constructed a SPLiCR-seq library that contains sgRNAs targeting all human RNA-binding proteins (RBPs) and transduced the library via lentivirus into HEK293T cells (CRISPRi-HEK293T) and iPSCs (CRISPRi-iPSC) that stably express CRISPRi machinery (dCas9-BFP-KRAB) from the CLYBL safe harbor locus. Following puromycin selection and cell expansion, we induced ER stress in these cells with Tg treatment for 3 hours (Fig. 2A). Subsequently, RNA was extracted and reverse transcribed, followed by PCR amplification of regions containing the splicing reporter and sgRNA sequences to generate NGS libraries for paired-end sequencing. From each read pair, Read2 identified the sgRNA while Read1 revealed the splicing outcome of the XBP1 reporter. Splicing phenotypes for each sgRNA were calculated as a ratio of spliced to unspliced read counts, normalized to non-targeting controls (Fig. 2A).

**Fig. 2:**
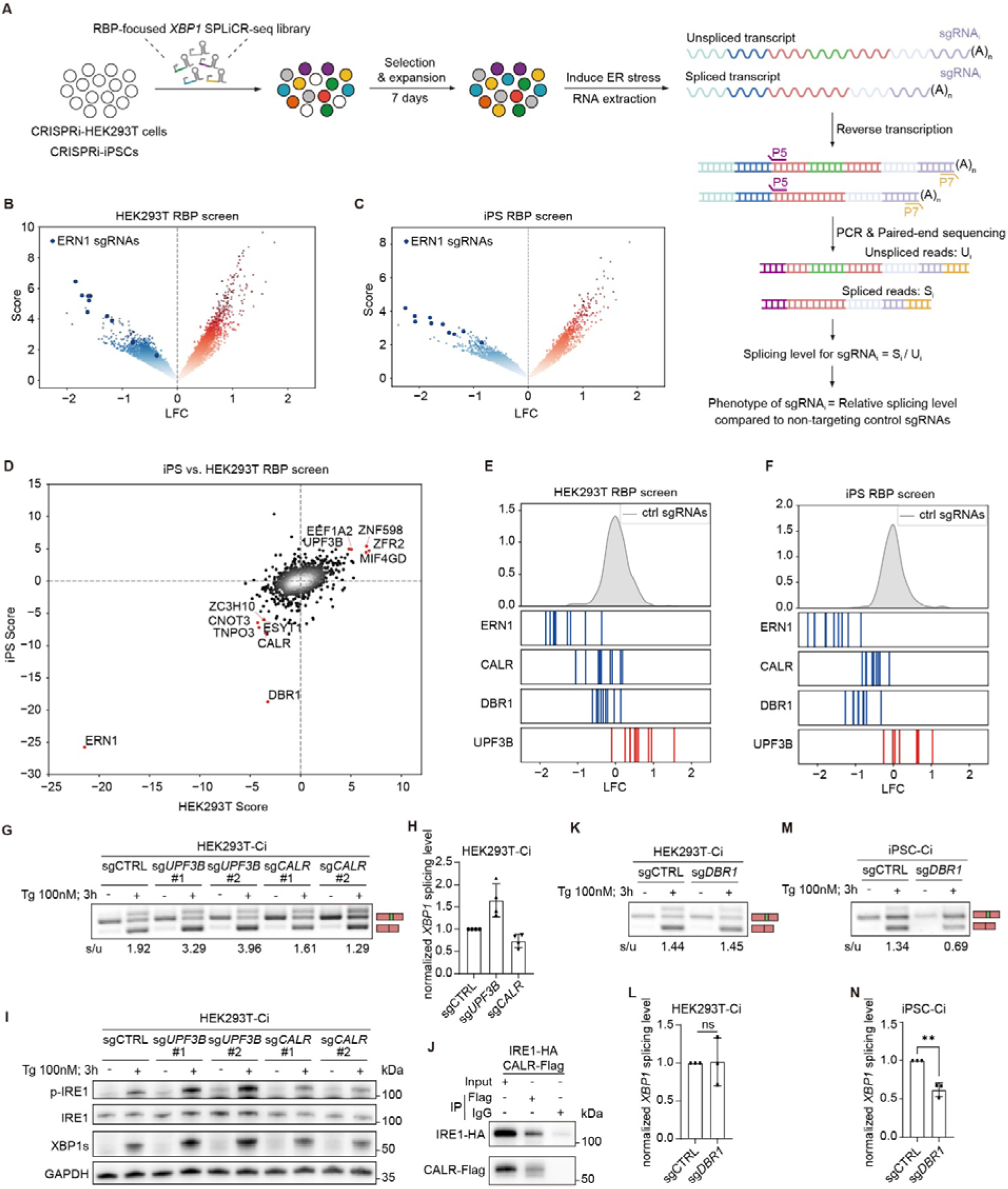
RBP-focused SPLiCR screens identify common and cell-type specific regulators of IRE-XBP1 Signaling. (A) Schematic illustration of the SPLiCR screen workflow. HEK293T cells and iPSCs expressing the CRISPRi machinery were transduced with an sgRNA library targeting 1,350 human RNA-binding proteins (MOI < 0.3). After puromycin selection and cell expansion, cells were treated with Tg for 3 hours to induce ER stress. RNA was extracted, reverse-transcribed, and regions containing the *XBP1* splicing reporter and sgRNA were PCR-amplified to generate paired-end NGS libraries. For each sgRNA, splicing phenotypes were calculated as the ratio of spliced reads (Si) to unspliced reads (Ui), and normalized to the median control sgRNA levels. The screen was performed in two biological replicates. **(B, C)** Volcano plots showing sgRNA phenotypes from the SPLiCR-seq screens in HEK293T cells and iPSC (**C**). sgRNAs targeting *ERN1* are highlighted in dark blue. **(I) (D)** Comparison of gene knockdown phenotypes from HEK293T and iPSC SPLiCR-seq screens, revealing both common and cell-type-specific regulators of *XBP1* splicing. **(E, F)** Distributions of phenotype scores for non-targeting control sgRNAs (gray) and sgRNAs targeting selected negative (blue) and positive (red) hits in HEK293T (**E**) and iPSC (**F**) SPLiCR-seq screens. **(G)** Representative RT-PCR analysis showing *XBP1* splicing in HEK293T cells following *CALR* or *UPF3B* knockdown with two independent sgRNAs. The ratios of spliced to unspliced *XBP1* (*XBP1s*/*XBP1u*) are indicated at the bottom. **(H)** Quantification of *XBP1* splicing levels for CALR and UPF3B knockdown in HEK293T cells, normalized to control sgRNA (mean ± SD, n = 4 technical replicates). **(I)** Western blot showing protein levels of XBP1s, IRE1 and phosphorylated IRE1 (p-IRE1) in HEK293T cells under ER stress conditions following knockdown of *CALR* or *UPF3B*. GAPDH was used as a loading control. **(J)** Co-immunoprecipitation showing a direct physical interaction between CALR and IRE1. **(K-N)** Representative RT-PCR analysis showing *XBP1* splicing under ER stress following *DBR1* knockdown in HEK293T cells (**K**) and iPSCs (**M**), with splicing levels indicated at the bottom. Quantification of RT-PCR results showing the effect of *DBR1* knockdown on *XBP1* splicing in HEK293T cells (**L**) and iPSCs (**N**). Splicing levels are normalized to control sgRNA levels (mean ± SD, n = 3 technical replicates). Student’s t-test: **p < 0.01; ns, not significant.

These screens identified genes whose knockdown either enhanced (positive hits) or suppressed (negative hits) *XBP1* splicing in both cell types (Fig. 2B, C, Supplementary Table 1). As anticipated, *ERN1* emerged as the top negative hit in both screens (Fig. 2B, C), validating our screening approach.

While most hits showed consistent effects across cell types, we also uncovered cell type-specific regulators (Fig. 2D). *DBR1,* for example, showed a strong negative phenotype in iPSCs but not in HEK293T cells (Fig. 2D). From the screen hits, we selected three candidates for validation: two negative regulators (*CALR* and *DBR1*) and one positive regulator (*UPF3B*) (Fig. 2E, F). *CALR* encodes calreticulin, an ER chaperone involved in protein folding and calcium homeostasis that was recently identified as an ATF6 repressor[30]. *DBR1* encodes an RNA debranching enzyme essential for intronic lariat degradation during splicing[38,39]. *UPF3B* is a component of the nonsense-mediated mRNA decay (NMD) pathway recently shown to interact with IRE1[40].

Validation experiments with two independent sgRNAs for each gene confirmed our screen results (Fig. 2G-N, Fig. S2). In HEK293T cells, knockdown of *UPF3B* significantly enhanced endogenous *XBP1* splicing (Fig. 2G, H), as well as XBP1s protein levels (Fig. 2I) under ER stress, while knockdown of *CALR* showed the opposite effect (Fig. 2G-I). Further investigation revealed that these effects were mediated through modulation of IRE1 activation, with *UPF3B* knockdown increasing and *CALR* knockdown decreasing IRE1 phosphorylation (Fig. 2I). In addition, our co-immunoprecipitation (co-IP) experiment revealed that CALR directly interacts with IRE1 (Fig. 2J). Notably, *DBR1* exhibited cell type-specific effects, with its knockdown reducing *XBP1* splicing specifically in iPSCs but not in HEK293T cells (Fig. 2K-N), consistent with the screen results and highlighting a cell-type-specific role for *DBR1* in modulating IRE1-XBP1 signaling. These results demonstrate the power of SPLiCR-seq to identify both common and cell type-specific regulators of RNA splicing across diverse cellular contexts.

### Genome-Wide SPLiCR-seq Screens Identify Regulators of IRE1-XBP1 Signaling Under Different ER Stress Conditions

Building on the success of the RBP-focused SPLiCR-seq screens, we expanded the platform to genome-wide screens to comprehensively identify regulators of IRE1-XBP1 signaling under different ER stress conditions. To this end, we constructed a genome-wide CRISPRi library containing more than 22,000 sgRNAs targeting over 11,000 genes into the XBP1 SPLiCR-seq vector. The screens were conducted following the same workflow as the RBP screen: CRISPRi-HEK293T cells were transduced with the library, treated with either Tg or Tm for 24 hours to induce ER stress, and RNA was extracted. SPLiCR libraries were then prepared for paired-end NGS to determine sgRNA-associated splicing phenotypes (Fig. 3A, Supplementary Table 1).

**Fig. 3:**
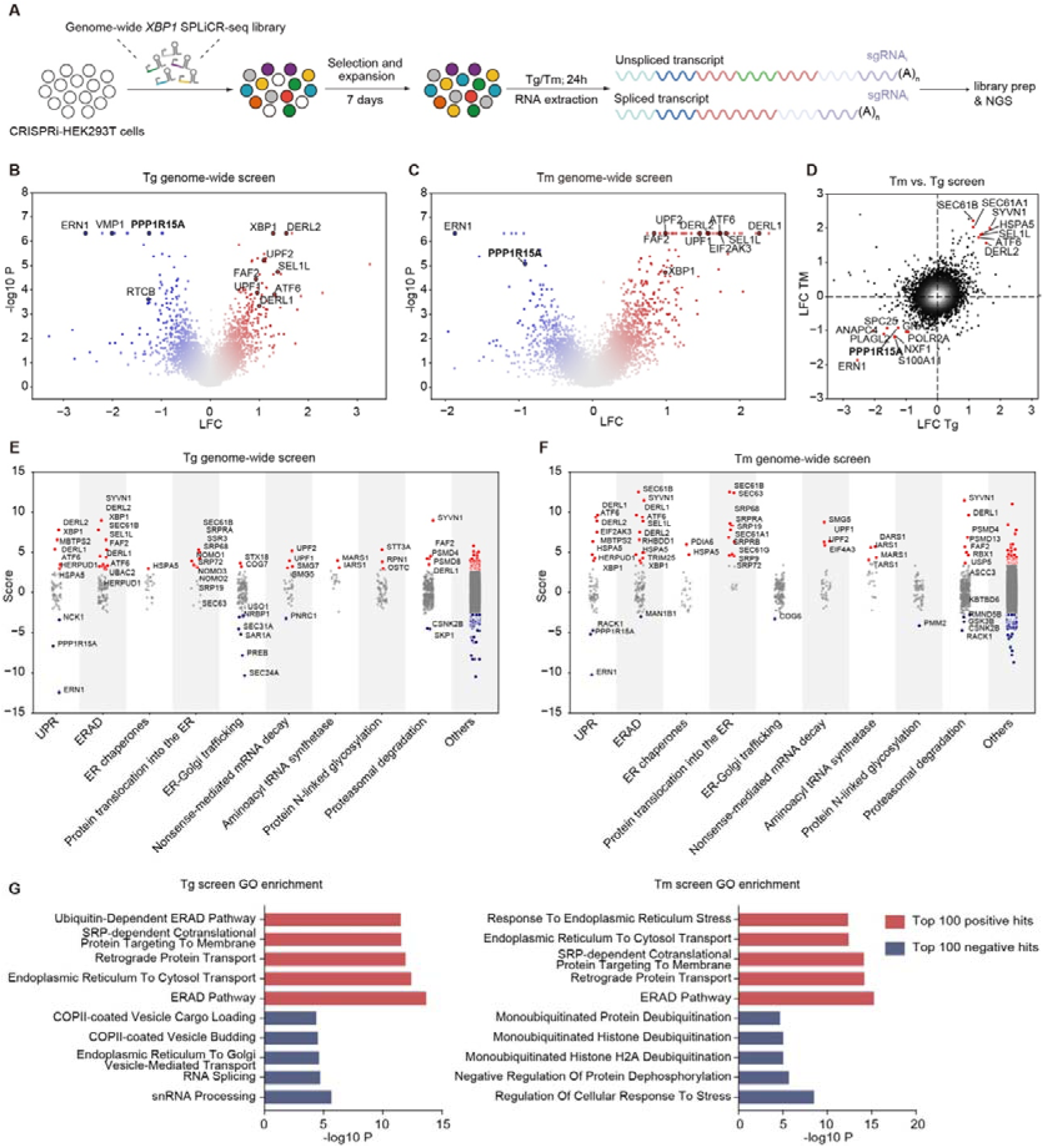
Genome-Wide SPLiCR-Seq Screens Identify Regulators of IRE1-XBP1 Signaling Under Different ER Stress Conditions. **(A)** Schematic of the genome-wide SPLiCR-seq screen workflow. CRISPRi-HEK293T cells were transduced with a genome-wide sgRNA library targeting 11,120 genes that are expressed in HEK293T cells (MOI < 0.3). Following puromycin selection and cell expansion, cells were treated with either Tg or Tm for 24 hours to induce ER stress. RNA was extracted and SPLiCR libraries were prepared for NGS. **(B, C)** Volcano plots showing gene knockdown phenotypes from genome-wide screens in HEK293T cells treated with Tg (**B**) or Tm (**C**). Key known regulators of the IRE1-XBP1 pathway are labeled, with *PPP1R15A*, a novel regulator investigated in this study, highlighted in bold. **(D)** Comparison of gene knockdown phenotypes between Tg and Tm screens, showing a strong overlap in common regulators of *XBP1* splicing as well as stressor-specific hits. **(E, F)** Functional categorization of hits from the genome-wide screens under Tg (**E**) and Tm (**F**) treatment. Red circles, positive hits; blue circles, negative hits. **(G)** GO Biological Process enrichment analysis of the top 100 positive hits (red) and top 100 negative hits (blue) for Tg (left) and Tm (right) treatments.

In both Tg and Tm screens, *ERN1* was identified as the top negative regulator, as expected, confirming the accuracy and robustness of the screens (Fig. 3B, C). Several known regulators of the IRE1-XBP1 pathway and broader UPR were also identified, including negative hits such as *VMP1*[41], *RTCB*[42], and *PPP1R15A*[43], and positive hits such as *UPF1*[40,44], *UPF2*[40,44], *DERL1* [45], *DERL2*[45], *ATF6*[6], *FAF2*[46], and *EIF2AK3*[36,37]. Additionally, a substantial fraction of our top hits overlaps with those discovered in a previous FACS-based genome-wide screen on IRE1-XBP1 signaling[47], further demonstrating the reliability and reproducibility of our SPLiCR-seq platform (Fig. S3).

A direct comparison of hits under Tg and Tm treatments revealed a strong overlap, with many genes showing consistent phenotypes across both conditions (Fig. 3D-G). Both screens identified regulators enriched in pathways associated with protein homeostasis and ER function, such as protein translation and degradation, UPR, ERAD, ER chaperones, and protein translocation into the ER (Fig. 3E-G). Importantly, the screens also identified stressor-specific regulators, indicating distinct mechanisms modulating IRE1-XBP1 signaling based on the type of ER stress. For instance, genes involved in COPII vesicle-mediated ER-to-Golgi trafficking, including *SEC24A*, *PREB*, *SAR1A*, and *SEC31A*, were significantly enriched among the top negative hits in the Tg screen but not the Tm screen (Fig. 3D-G). In contrast, pathways related to post-translational modification were uniquely enriched in the Tm screen, underscoring the distinct cellular responses to these two stressors.

These genome-wide SPLiCR-seq screens provide a comprehensive catalog of regulators involved in IRE1-XBP1 signaling and UPR, uncovering both conserved and stressor-specific regulatory mechanisms. Together, these results establish SPLiCR-seq as a robust and scalable platform for probing RNA splicing.

### Batch validation screens identify *PPP1R15A*(GADD34) as a novel regulator of IRE1-XBP1 signaling

To validate the hits from our genome-wide screens, we conducted a secondary batch validation screen using a targeted library containing sgRNAs against 70 selected hits. This validation screen followed the same workflow as the primary screen (Fig. 4A). Most genes in the validation screen exhibited knockdown phenotypes consistent with their effects observed in the primary screens, confirming the reliability of our findings (Fig. 4B, Supplementary Table 1).

**Fig. 4:**
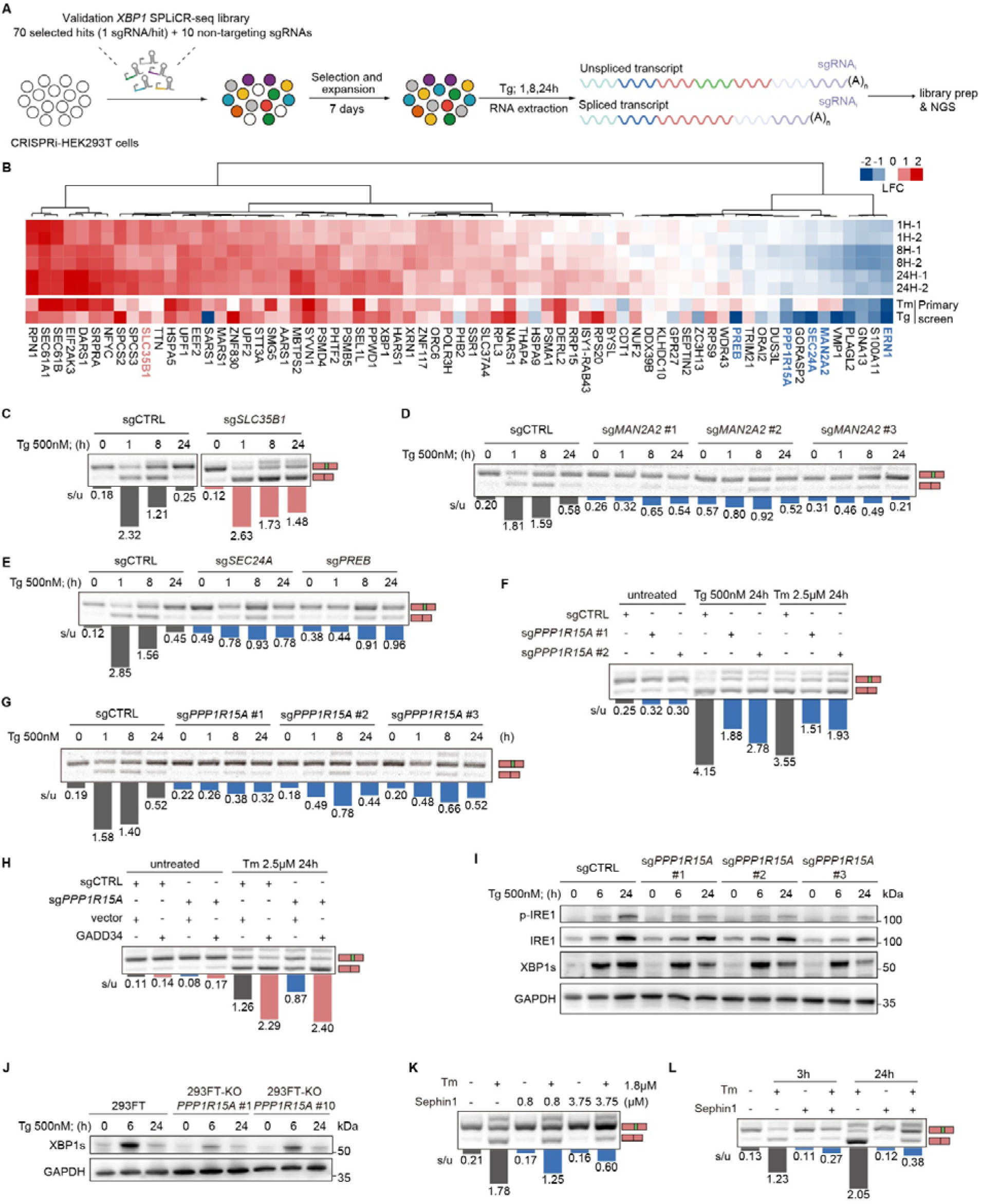
Validation Screens Identify PPP1R15A (GADD34) as a Novel Regulator of IRE1-XBP1 Signaling. **(A)** Schematic of the batch validation SPLiCR-seq screen workflow. **(B)** Heatmap showing gene knockdown phenotypes from the validation screens (top), compared to the primary screens (bottom). Genes are hierarchical clustered based on their validation screen phenotypes. Genes selected for individual validation are highlighted in blue (negative hits) and red (positive hits). **(C-F)** RT-PCR validations of *XBP1* splicing in HEK293T cells under Tg- or Tm-induced ER stress following knockdown of *SLC35B1* (**C**), *MAN2A2* (**D**), *SEC24A*(**E**), *PREB* (**E**) and *PPP1R15A* (**F**). Splicing ratios are indicated below each lane. **(G)** Time-course RT-PCR analysis showing *XBP1* splicing in *PPP1R15A* knockdown cells treated with Tg (500 nM) for the indicated durations. Splicing ratios are indicated below each lane. **(H)** RT-PCR analysis of *XBP1* splicing in control and *PPP1R15A* knockdown HEK293T cells expressing either empty vector or GADD34 vector. Splicing ratios are indicated below each lane. **(I)** Western blot analysis of p-IRE1, total IRE1, and XBP1s protein levels in control and *PPP1R15A* knockdown cells. GAPDH was used as a loading control. **(J)** Western blot analysis of XBP1s protein levels in control and *PPP1R15A* knockout cells. GAPDH was used as a loading control. **(K, L)** RT-PCR analysis of *XBP1* splicing in HEK293T cells treated with varying concentrations of Sephin1 (**K**) and at varying durations of Tm treatment (**L**). Splicing ratios are indicated below each lane.

From these high-confidence hits, we selected several genes for individual validation of their effects on endogenous *XBP1* splicing in UPR. These included the positive hit *SLC35B1*, which encodes a nucleotide sugar transporter required for protein glycosylation[48,49], and negative hits *MAN2A2*, *SEC24A*, *PREB*, and *PPP1R15A*. Among these, MAN2A2 regulates N-glycan processing[50], *SEC24A* and *PREB* are involved in the COPII vesicle-mediated ER-Golgi trafficking pathway[51,52], and *PPP1R15A* encodes GADD34, known to promote ER stress recovery through eIF2α dephosphorylation[43,53,54].

Consistent with the screen results, knockdown of *SLC25B1* dramatically enhanced endogenous *XBP1* splicing at all Tg treatmemt time-1, 8 and 24 hrs, suggesting enhanced IRE1-XBP1 signaling (Fig. 4C, Fig. S4A). Conversely, knockdown of *MAN2A2*, *SEC24A*, *PREB*, *PPP1R15A* significantly suppressed *XBP1* splicing (Fig. 4D-G, Fig. S4A-E).

Given its strong phenotypes in both Tm and Tg screens and previously uncharacterized role in IRE1-XBP1 signaling, we further investigated *PPP1R15A* (GADD34). Overexpression of GADD34 in *PPP1R15A* (GADD34) knockdown cells rescued the *XBP1* splicing phenotype, while overexpression in wild-type cells further enhanced *XBP1* splicing (Fig. 4H, Fig. S4F). At the protein level, western blot analysis confirmed that both knockdown and knockout of *PPP1R15A* reduced XBP1s protein levels under ER stress (Fig. 4I, J, Fig. S4G-I). Additionally, both knockdown and knockout of *PPP1R15A* suppressed IRE1 activation, as evidenced by reduced phosphorylated IRE1 (p-IRE1) levels (Fig. 4I, Fig. S4I). These results suggest that GADD34 regulates *XBP1* splicing by modulating IRE1 activation.

To complement our genetic approaches, we examined the effects of pharmacological GADD34 inhibition using Sephin1, a small-molecule inhibitor of GADD34[55]. Similar to genetic knockdown, Sephin1 treatment potently suppressed *XBP1* splicing in a dose-dependent manner and across different ER stress treatment durations (Fig. 4K, L).

Together, these results confirm the discovery of novel IRE1-XBP1 signaling regulators from the genome-wide screens and establish GADD34 as a previously unrecognized modulator of this pathway.

### GADD34 Regulates IRE1-XBP1 Signaling Through an eIF2**α**-Independent Mechanism

GADD34 has previously been proposed to function as a negative feedback regulator of the UPR by recruiting Protein Phosphatase 1 (PP1) to dephosphorylate eIF2α[56–58], which is phosphorylated by PERK during ER stress. However, our findings reveal that GADD34 regulates IRE1-XBP1 signaling through a mechanism independent of eIF2α, supported by several lines of evidence:

First, we found no increase in eIF2α phosphorylation levels in HEK293T cells subjected to either genetic or pharmacological inhibition of GADD34 under Tg- or Tm-induced ER stress conditions (Fig. 5A, B, S4I, S5A). Moreover, the ability of GADD34 to regulate eIF2α phosphorylation was dependent on its expression levels: notable dephosphorylation of eIF2α was only observed with massive GADD34 overexpression achieved through transient transfection, while moderate overexpression via lentiviral infection produced no notable changes in eIF2α phosphorylation (Fig. 5C).

**Fig. 5:**
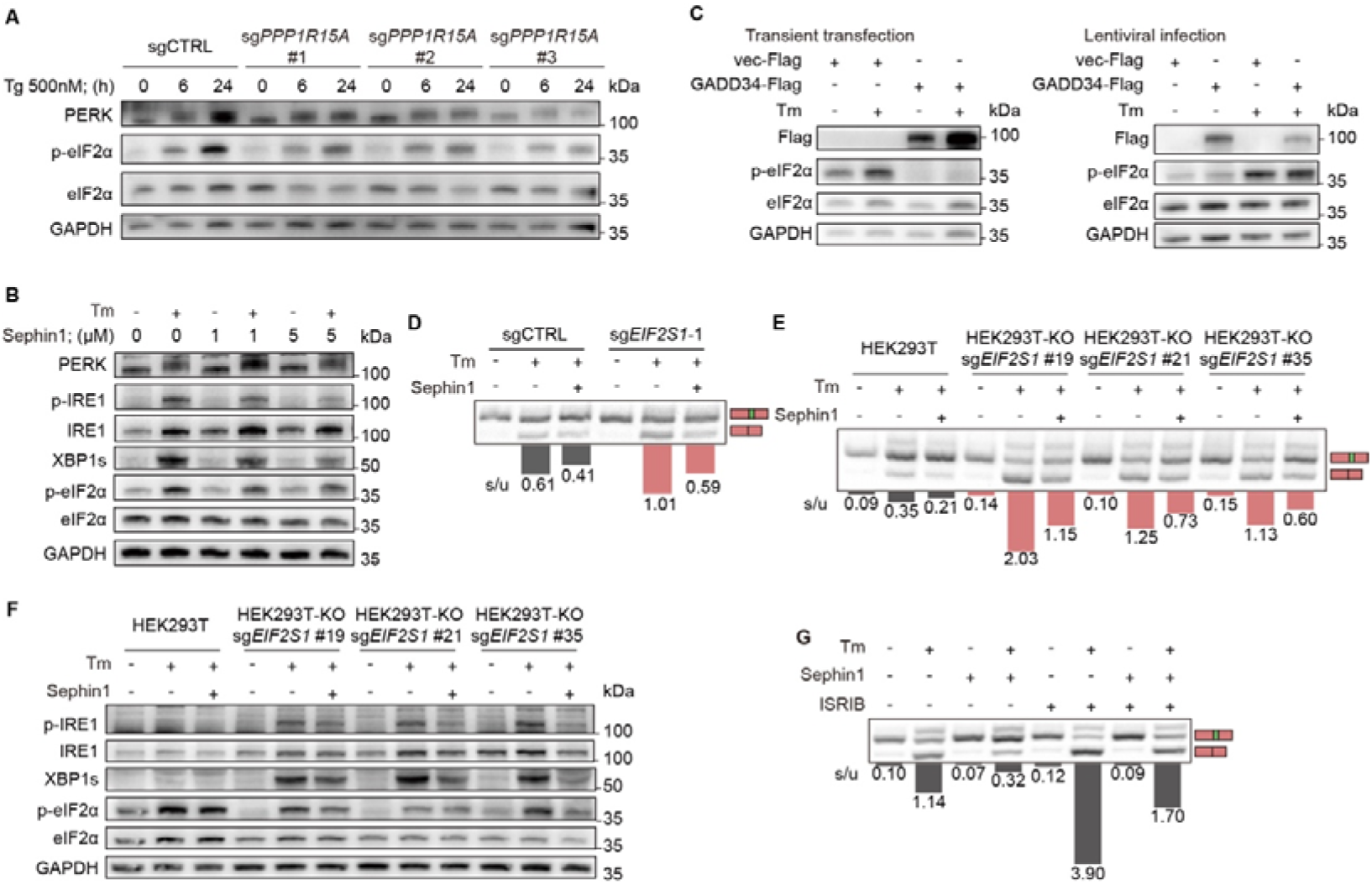
GADD34 Regulates IRE1-XBP1 Signaling Through an eIF2α-Independent Mechanism. **(A)** Western blot analysis of PERK, phosphorylated eIF2α (p-eIF2α) and total eIF2α levels in control and *PPP1R15A* knockdown HEK293T cells treated with Tg (500 nM) for the indicated durations (0, 6, or 24 hours). GAPDH was used as a loading control. **(B)** Western blot analysis of PERK, p-eIF2α, total eIF2α, p-IRE1, total IRE1, XBP1s and ATF4 in HEK293T cells treated with increasing concentrations of Sephin1 (0, 1, 5, or 10 μM) in the presence or absence of Tm (2 μM, 24 hours). GAPDH was used as a loading control. **(C)** Western blot analysis of p-eIF2α levels in cells with GADD34 overexpression via transient transfection (left) or lentiviral infection (right). GAPDH was used as a loading control. **(D, E)** RT-PCR analysis of *XBP1* splicing in control and *EIF2S1* (eIF2α) knockdown cells (**D**), and *EIF2S1* partial knockout cells (**E**) treated with Tm (2 μM, 24 h) and Sephin1 (5 μM, 24 h) as indicated. Splicing ratios are indicated below each lane. **(F)** Western blot analysis of p-eIF2α, total eIF2α, p-IRE1, total IRE1 and XBP1s in control and *EIF2S1* partial knockout HEK293T cells treated with Tm (2 μM, 24 h) and Sephin1 (5 μM, 24 h) as indicated. GAPDH was used as a loading control. **(G)** RT-PCR analysis of *XBP1* splicing in HEK293T cells treated with Tm (2 μM, 24 h), Sephin1 (5 μM, 24 h), and ISRIB (1 μM, added 3 h before collection) as indicated. Splicing ratios are indicated below each lane.

Second, we generated *eIF2*α knockdown and partial knockout HEK293T cell lines (complete knockout of eIF2α was found to be lethal) (Fig. S5B, C). These cells exhibited enhanced UPR activation under ER stress, as evidenced by increased IRE1 phosphorylation, XBP1 splicing, and ATF4 expression. Notably, despite the elevated levels of UPR activation, GADD34 inhibition by Sephin1 remained effective in suppressing IRE1-XBP1 signaling in these eIF2α-depleted cells, as evidenced by reduced *XBP1* splicing, XBP1s expression and IRE1 phosphorylation (Fig. 5D-F).

Finally, treatment with ISRIB, a small-molecule activator of eIF2B that reverses the translational inhibition caused by eIF2α phosphorylation[59], failed to prevent Sephin1’s inhibitory effect on XBP1 splicing during ER stress (Fig. 5G, S5A).

These results collectively demonstrate that GADD34 regulates IRE1-XBP1 signaling through an eIF2α-independent mechanism.

### GADD34 Regulates IRE1-XBP1 Signaling Through Direct Interaction with IRE1

To investigate how GADD34 regulates IRE1-XBP1 signaling, we examined whether GADD34 directly interacts with IRE1. Co-IP experiments revealed that full-length GADD34 (GADD34 FL) physically interacts with IRE1 under both basal and ER stress conditions (Fig. 6A, B).

**Fig. 6:**
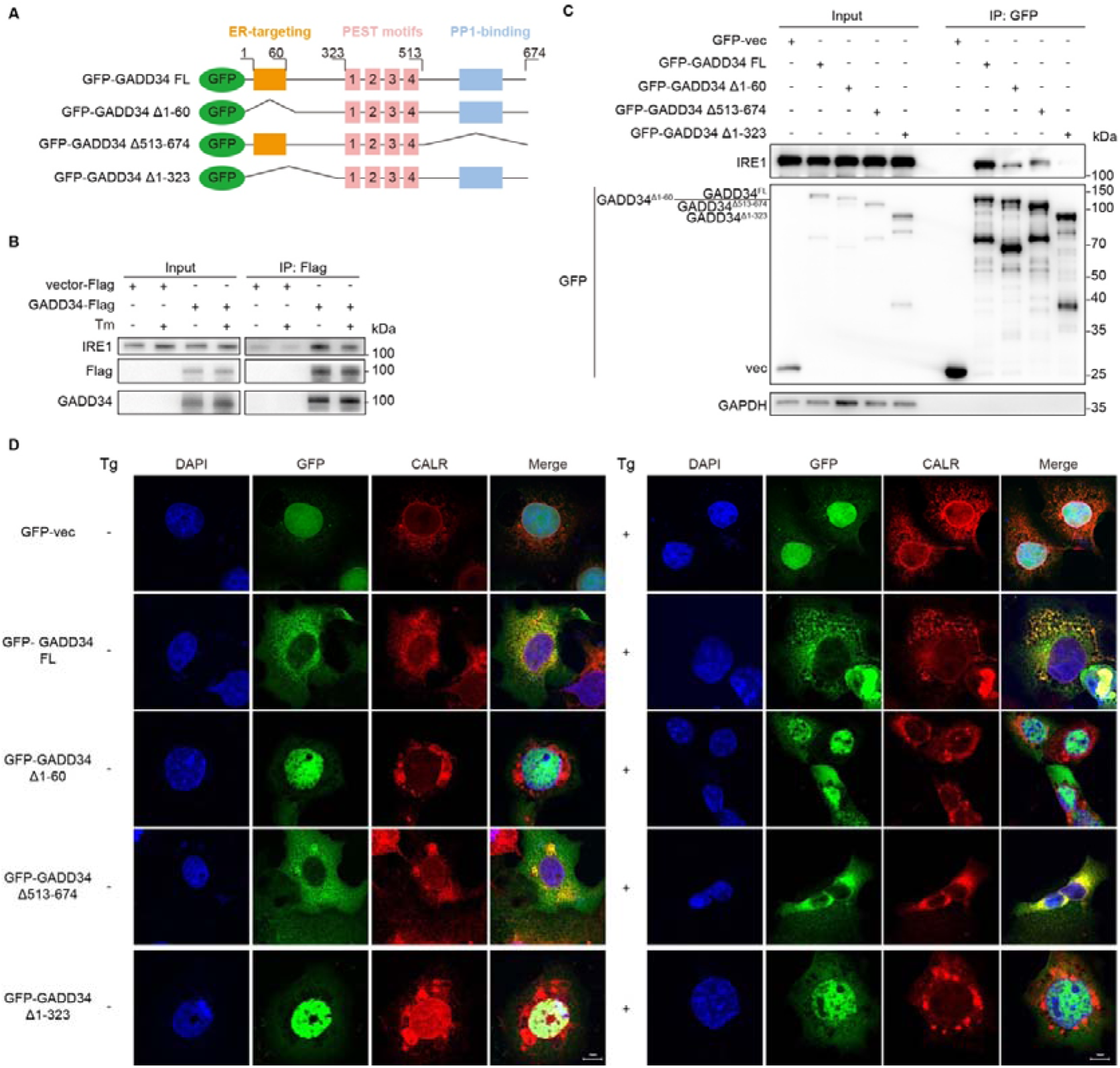
GADD34 Directly Interacts with IRE1. **(A)** Schematic representation of full-length GADD34 (GADD34 FL) and its truncation mutants analyzed in the study. The ER-targeting domain (residues 1-60), PEST motifs (residues 323-513), and PP1-binding domain (residues 513-674) are indicated. GFP was fused to the N-terminus of each construct. **(B, C)** Co-IP analysis of the interaction between IRE1 and full-length GADD34 (**B**) or truncated GADD34 variants(**C**). **(D)** Immunofluorescence showing subcellular localization of GADD34 variants. HEK293T cells expressing GFP-vector or indicated GFP-GADD34 constructs (green) were treated with or without Tg (2 μM, 3h) and immunostained for the ER marker calreticulin (CALR, red). Nuclei were counterstained with DAPI (blue). Scale bar = 10 μm.

GADD34 contains a putative ER-targeting domain at its N-terminus, four PEST repeats, and a PP1-binding domain at the C-terminus [60](Fig. 6A). To determine which domains are crucial for IRE1 binding, we generated several GADD34 truncation mutants (Fig. 6A): a large N-terminal deletion (Δ1–323), a smaller N-terminal deletion that removes the putative ER-targeting domain (Δ1–60), and a C-terminal deletion that removes the PP1-binding domain (Δ513-674). Co-IP experiments revealed that deletion of the ER-targeting domain (GADD34 Δ1-60) or the PP1-binding domain (GADD34 Δ513-674) greatly reduced the GADD34-IRE1 interaction, while the large N-terminal deletion (GADD34 Δ1-323) completely abolished the interaction (Fig. 6C). These results suggest that the N-terminal region, specifically the ER-targeting domain, and the PP1-binding domain play a critical role in mediating the interaction between GADD34 and IRE1.

Confocal microscopy analysis showed that GADD34 FL predominantly localizes to the ER, as evidenced by co-localization with the ER marker calreticulin (CALR), under both basal and Tg-induced ER stress conditions (Fig. 6D). However, both N-terminal truncations (GADD34 Δ1-323 and GADD34 Δ1-60) disrupted this ER localization and predominantly accumulated in the nucleus. In contrast, deletion of the PP1-binding domain (GADD34 Δ513-674) did not notably affect GADD34’s ER localization (Fig. 6D).

Together, these results indicate that GADD34 directly interacts with IRE1 and that its N-terminal region, particularly residues 1–60, is crucial for both ER localization and robust binding to IRE1. Although the PP1-binding domain does not determine GADD34’s subcellular localization, it contributes to the overall stability or efficiency of the GADD34–IRE1 interaction.

### GADD34 Inhibition by Sephin1 Mitigates CAR-T Cell Exhaustion and Enhances Tumor Killing

Previous studies have implicated ER stress and XBP1 splicing in driving T cell functional exhaustion and tumor immunoevasion[15,16,61]. Hence, we speculated that GADD34 inhibition may delay or reverse T cell exhaustion, leading to enhanced anti-tumor activity. To model T cell exhaustion in vitro, we leveraged a serial co-culture assay where human chimeric antigen receptor T cells (CAR-T cells) were repetitively stimulated with target tumor cells. CAR-T cells exhibit gradual loss of effector functions and reduction in anti-tumor capacity upon each round of co-culture, indicative of functional exhaustion[62].

To test our hypothesis, B7-H3 specific CAR-T cells were co-cultured with pancreatic cell line BXPC3 in the presence of DMSO or GADD34 inhibitor Sephin1 (Fig. 7A). We observed significant increase in *XBP1* splicing in CAR-T cells after 3 rounds of stimulation (D9), which was effectively suppressed upon Sephin1 addition (Fig. 7B-C). Importantly, CAR-T cells treated with Sephin1 maintained superior effector functions, as indicated by sustained IFN-γ and TNF-α secretion, especially after the second and third rounds of co-culture (Fig. 7D-G).

**Fig. 7:**
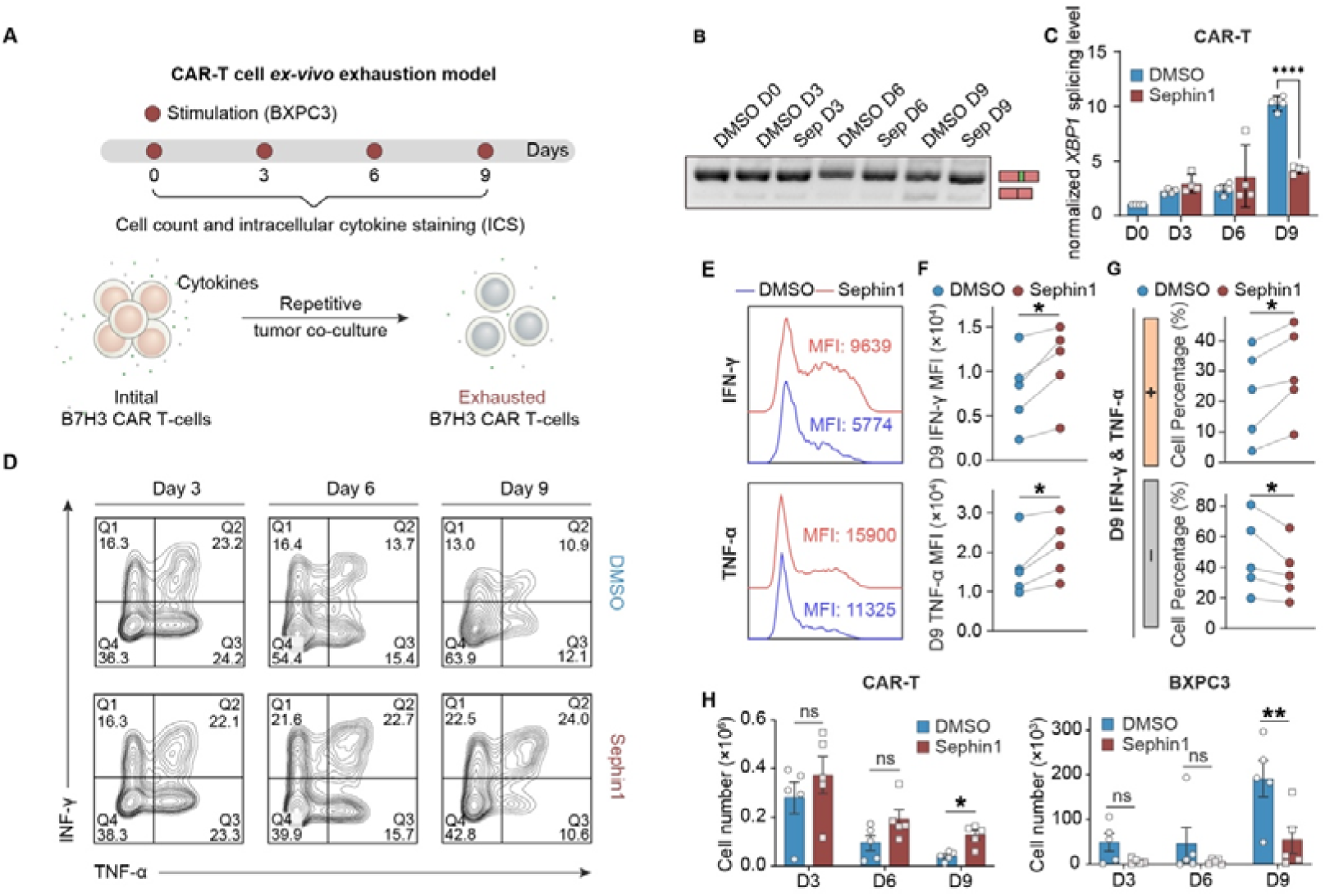
Sephin1 reduces XBP1 splicing in *ex-vivo* exhaustion model thereby alleviating CAR-T cell exhaustion. **(A)** Schematic diagram of the *ex-vivo* CAR-T cell exhaustion model. **(B)** Representative agarose gel showing RT-PCR products of endogenous XBP1 splicing in CAR-T cells repetitively co-cultured with tumor cells for 3, 6 and 9 days in the presence of DMSO or Sephin1. **(C)** *XBP1* splicing level in CAR-T cells repetitively co-cultured with tumor cells for 3, 6 and 9 days in the presence of DMSO or Sephin1 and normalized to that of the DMSO treatment in day 0 and presented as fold change. Data are presented as mean ± SD, paired Student’s t-test; ***P<0.0001, n = 4 technical replicates). **(D, E)** Intracellular cytokine staining of IFN-γ and TNF-α in CAR-T cells repetitively co-cultured with tumor cells for 3, 6 and 9 days in the presence of DMSO or Sephin1 and the percentage of IFN-γ^+^/TNF-α^+^ and IFN-γ^-^/TNF-α^-^ in CAR-T cells at day 9. (Data are presented as mean ± SEM, paired Student’s t-test; *P < 0.05, n = 5 experiments). **(F, G)** Expression of of IFN-γ and TNF-α in CAR-T cells at day 9 and the mean fluorescent intensity (MFI) of IFN-γ and TNF-α in cells stained positive for respective cytokine. (Data are presented as mean ± SEM, paired Student’s t-test; *P < 0.05, n = 5 experiments). **(H)** Cell counts of CAR-T cells and BXPC3 cells at day 3, 6, 9 upon treatment of DMSO or Sephin1 (Data are presented as mean ± SEM, two-way ANOVA with Sidak multiple comparisons test; ns, not significant, *P < 0.05, **P < 0.01, n = 5 experiments).

Furthermore, Sephin1 treatment enhanced both the expansion and anti-tumor activity of CAR-T cells, as evidenced by increased numbers of CAR-T cells and reduced residual tumor cells after each round of co-culture (Fig. 7H). To rule out potential direct effects of Sephin1 on cell viability, we confirmed that addition of Sephin1 to tumor or CAR-T cells alone does not alter their cell numbers (Fig. S6A, B).

Notably, the protective effect of Sephin1 was not limited to the B7-H3 CAR-T/BXPC3 model. We observed similar improvements when testing CD19-specific CAR-T cells co-cultured with breast cancer cell line HCC1806 expressing the CD19 antigen (Fig. S6C-G), suggesting that the beneficial effects of GADD34 inhibition on CAR-T cell function are not limited to specific antigens or tumor models.

Collectively, these findings demonstrate that GADD34 inhibition by Sephin1 suppresses *XBP1* splicing and alleviates CAR-T cell exhaustion, leading to enhanced CAR-T cell expansion, sustained effector function, and improved tumor killing. These results highlight the therapeutic potential of GADD34 inhibition for improving CAR-T tumor immunotherapy.

## Discussion

In this study, we developed SPLiCR-seq, a novel CRISPR-based screening platform for studying RNA splicing regulation (Fig. 1A). Unlike existing screening methods that rely on indirect fluorescence-based readouts and FACS-based sorting[18–30], SPLiCR-seq links genetic perturbations to direct readout of RNA splicing phenotypes via NGS. This approach provides several key advantages: it enables high-resolution analysis of splicing dynamics with improved temporal precision, eliminates the lag between splicing events and readout signals inherent to fluorescent reporters, and extends applicability to diverse biological contexts where FACS-based sorting is impractical. Our validation experiments confirmed that the SPLiCR-seq vector accurately reflects endogenous splicing dynamics, as demonstrated by the dose- (Fig. 1D-E) and time-dependent (Fig. 1F-H, Fig. S1A, B) splicing of *XBP1* under ER stress.

The power of SPLiCR-seq was demonstrated through our targeted and genome-wide screens, which revealed both conserved and context-specific regulators of *XBP1* splicing under ER stress. Our RBP-focused screens across different cell types uncovered cell type-specific regulation of splicing, exemplified by *DBR1*, which showed strong phenotypes specifically in iPSCs but not in HEK293T cells (Fig. 2K-N). This finding highlights the importance of cellular context in splicing regulation and demonstrates SPLiCR-seq’s utility in uncovering context-dependent mechanisms. The genome-wide screens further expanded our understanding of IRE1-XBP1 regulation, identifying both conserved regulators across different ER stressors and stressor-specific modulators. For instance, genes involved in COPII vesicle-mediated ER-to-Golgi trafficking were specifically enriched among negative regulators under Tg treatment but not Tm (Fig. 3G), revealing distinct regulatory mechanisms engaged by different types of ER stress.

Our screens yielded comprehensive catalogs of *XBP1* splicing regulators across diverse contexts, providing valuable insights into both IRE1-XBP1 signaling and broader UPR regulation. Among the identified regulators, PPP1R15A (GADD34) emerged as a particularly intriguing hit due to its strong and consistent phenotypes across multiple conditions (Fig. 3B-C). GADD34 has been previously established as a negative feedback regulator of the unfolded protein response (UPR)[63], functioning to dephosphorylate eIF2α by recruiting protein phosphatase 1 (PP1). The small molecule inhibitor Sephin1 (also known as Icerguastat) was first reported by Das et al.[55] as a selective inhibitor of GADD34 that disrupts the GADD34-PP1 interaction. Sephin1 has demonstrated therapeutic potential in several diseases, including Charcot-Marie-Tooth disease type 1B[55], amyotrophic lateral sclerosis (ALS)[55], multiple sclerosis[64], and spinocerebellar ataxia[65]. It is currently undergoing Phase 2 clinical trials for ALS. However, the proposed mechanism of action of Sephin1 has been contested. Crespillo-Casado et al.[66] found that Sephin1 did not disrupt the formation or stability of the GADD34-PP1 complex in vitro biochemical assays and had no measurable effect on eIF2α dephosphorylation in cells.

In our study, we observed that neither genetic inhibition of GADD34 via CRISPR nor pharmacological inhibition using Sephin1 resulted in increased phosphorylation levels of eIF2α in HEK293T cells (Fig. 5A, B, S4I, S5A). Interestingly, significant eIF2α dephosphorylation was only observed when GADD34 was massively overexpressed in cells using transient transfection, but not when expressed at more physiological levels via lentiviral infection (Fig. 5C). This raises the possibility that GADD34’s canonical role in eIF2α dephosphorylation may be context-dependent or secondary to other functions, particularly under conditions of supraphysiological expression.

Instead of affecting eIF2α phosphorylation, we found that GADD34 inhibition strongly reduced *XBP1* splicing under ER stress (Fig. 4F-G). Further analysis revealed that GADD34 modulates IRE1 activation and directly interacts with IRE1 (Fig. 6B). This interaction likely involves multiple regions of GADD34, including its ER-targeting region and PP1-binding domain. These findings suggest that GADD34 plays a previously unrecognized role in regulating IRE1-XBP1 signaling, independent of its canonical function in eIF2α dephosphorylation. The precise molecular mechanism by which GADD34 modulates IRE1 activation remains to be elucidated.

Building on these findings, we explored the therapeutic potential of targeting GADD34 in cancer immunotherapy. ER stress and *XBP1* splicing have been implicated in T cell exhaustion, a major barrier to the efficacy of CAR-T cell immunotherapy[12,15,16,61]. By inhibiting GADD34 with Sephin1, we were able to suppress *XBP1* splicing and alleviate CAR-T cell exhaustion in an *ex vivo* serial co-culture model (Fig. 7A). Sephin1-treated CAR-T cells exhibited enhanced expansion, sustained effector function, and improved tumor-killing capacity across multiple cancer models, including both B7-H3 (Fig. 7D-H) and CD19 CAR-T cells (Fig. S6C-G), suggesting that GADD34 inhibition may represent a broadly applicable strategy for improving CAR-T cell performance. Future studies should focus on evaluating the *in vivo* efficacy and safety profile of GADD34 inhibition to fully assess its potential as a therapeutic strategy for improving CAR-T cell performance.

The modular "plug-and-play" design of our SPLiCR-seq platform provides broad applicability for investigating a wide array of splicing events. While we demonstrated its utility in dissecting *XBP1* splicing regulation, the system can be easily adapted to investigate other critical splicing events, in particular those implicated in human diseases, such as SMN2 splicing in spinal muscular atrophy[67], *LMNA* splicing in progeria[67], cryptic splicing in TDP-43 proteinopathies[4] and *MAPT* splicing in Alzheimer’s disease and frontotemporal dementia[4]. By systematically identifying the key regulators of these splicing events, SPLiCR-seq holds the potential to advance our understanding of RNA splicing regulation and facilitate the development of novel therapeutic strategies against splicing-related diseases.

## Materials and Methods

### Cell culture

HEK293T and 293FT cells were purchased from ATCC and cultured in Dulbecco’s Modified Eagle’s Medium (Gibco, C11995500BT) supplemented with 10 % fetal bovine serum (TransGen Biotech, FS301-02) and 1 % penicillin-streptomycin (Aladdin, P301861). iPSCs (WTC11 background) were cultured in StemFlex medium (Gibco, A3349401) and plated on Matrigel (Corning, 356231) coated plates for at least 30 minutes before seeding the cells. 10 nM ROCK inhibitor (Selleck, S1049) was added on the first day of plating iPSCs. Human PDAC cell line, BXPC3 and TNBC cell line, HCC1806, were purchased from Procell (Wuhan, China) and were maintained in RPMI1640 (Gibco, C11875500BT) supplemented with 10% FBS (TransGen, FS301-02), 2 mM GlutaMax (Thermo, 35050061) and 100 U/mL penicillin and 100 μg/mL streptomycin. Human T cells were maintained in complete human T cell medium (1/2 RPMI-1640, 1/2 Click’s Media (Mesgen, MCM4125), supplemented with 10% FBS (Hyclone, SV30208.02), 2mM GlutaMAX, 100 U/mL of Penicillin and 100 μg/mL of streptomycin). Expression of surface markers and functional readouts were validated by FACS.Cells. All cells were maintained at 37°C and 5% CO2, and were tested negative for mycoplasma contamination using the MycAway™ Plus-Color One-Step Mycoplasma Detection Kit (Yeasen, 40612ES25).

### Generation of the XBP1 SPLiCR-seq vector

Sequences flanking the splicing site of *XBP1* was amplified from HEK293T cDNA by PCR. The amplified product was inserted into the pMK1334 vector by replacing the original WPRE cassette between EcoRI and SalI sites using the ClonExpress® Ultra One Step Cloning Kit (Vazyme, C115). pMK1334 was a gift from Martin Kampmann (Addgene plasmid # 127965 ; http://n2t.net/addgene:127965 ; RRID:Addgene_127965).

### sgRNA cloning

sgRNAs for CRISPRi were cloned into the SPLiCR-seq vector or pLG15 via BstXI and Bpu1102I sites as previously described[31]. sgRNAs for CRISPR knockout were cloned into the pX459 vector via BsmBI sites. The pX459 vector was a gift from Feng Zhang (Addgene plasmid # 62988 ; http://n2t.net/addgene:62988 ; RRID:Addgene_62988). A complete list of sgRNA sequences used in this study is listed in Supplementary Table 2.

### Lentivirus production

For lentivirus production of the sgRNA library, 5 × 10L HEK293T cells were seeded in a 15 cm dish for 24 hrs before transfection. Fifteen micrograms of sgRNA library plasmid and 15 µg of third-generation packaging mix (1:1:1 mix of the three plasmids: pMDLg/pRRE (Addgene, 12251), pRSV-Rev (Addgene, 12253), and pMD2.G (Addgene, 12259)) were diluted in 3 mL of Opti-MEM (Gibco, 31986-07). Subsequently, 45 µL of 2 mg/mL Polyethylenimine Linear (PEI) MW40000 (Yeasen, 40816ES03) was added to the 3 mL DNA dilution, vortexed for 10 seconds, and thoroughly mixed. After incubating at room temperature for 15-20 minutes, the mixture was added to the 15 cm dish containing HEK293T cells. 48 hrs later, the viral supernatants were collected and filtered through a 0.45 μm filter (Millipore, SLHV033RB).

For small-scale lentivirus production, 0.5 × 10L HEK293T cells were seeded on 6-well plates. After 24 hours, 100 µL of Opti-MEM was used to dilute 1 µg of transfer plasmid and 1 µg of third-generation packaging mix, thoroughly mixed to form the DNA dilution. Subsequently, 3 µL of PEI was added, and the mixture was vortexed for homogeneity. The remaining procedures were carried out as described above.

### Quantitative real-time polymerase chain reaction (qPCR)

Total RNA was extracted using the MolPure® Cell RNA Kit (Yeasen, 19231ES50) according to the manufacturer’s instructions. RNA was reverse transcribed to cDNA with the TransScript® One-Step gDNA Removal and cDNA Synthesis SuperMix (TransGen, AT311-03). Quantitative real-time PCR was performed using AceQ qPCR SYBR Green Master Mix (Vazyme, CQ111-02) according to the manufacturer’s protocol and run on a Fluorescence Quantitative PCR detection system (FDQ-96A). GAPDH was used as an endogenous control.

### Detection of *XBP1* splicing in SPLiCR vector by RT-PCR and Grayscale analysis

To measure RNA editing levels in the SPLiCR vector, cDNA products were used as templates for PCR amplification of fragments flanking the *XBP1* site. The PCR products were loaded onto a 3% agarose gel, imaged using a fully automated gel imaging system (OI 1000), and then analyzed for grayscale intensity in Fiji.

### CRISPRi screening with SPLiCR-seq and data analysis

The human RNA-binding protein (RBP) sgRNA library (6,853 unique sgRNA sequences targeting 1,350 RBPs, along with 250 non-targeting control sgRNAs) and the genome-wide HEK293T sgRNA library (22,240 unique sgRNA sequences targeting 11,120 genes expressed in HEK293T, along with 1,760 non-targeting control sgRNAs) were synthesized by GENEWIZ and cloned into the XBP1 SPLiCR-seq vector between BstXI and BlpI sites. To evaluate the library quality, the fragment harboring sgRNA sequences was amplified using Phanta Flash Master Mix (Vazyme, P520) following the manufacturer’s instructions, and the PCR products were analyzed by next-generation sequencing.

Lentivirus production of the SPLiCR-seq library was carried out as described above. The lentivirus was transduced into CRISPRi-HEK293T and CRISPRi-iPSC cells at a multiplicity of infection (MOI) less than 0.3. 48 hours later, the transduced cells were selected with 2 µg/mL puromycin for 48 hours to eliminate uninfected cells and generate a genome-edited cell pool. After 7 days of passage in medium without puromycin and treated with Thapsigargin (Tg, Santa Cruz, sc-24017) or Tunicamycin (Tm, Beyotime, SC0393), 10 million cells were collected for total RNA extraction using the MolPure® Cell RNA Kit. The mRNA with a poly-A tail was purified from the total extracted RNA using the BeyoMag™ mRNA Purification Kit with Magnetic Beads (Beyotime, R0071M) and reverse transcribed into cDNA using Oligo-dT primers and TransScript® II One-Step gDNA Removal and cDNA Synthesis SuperMix (TransGen, AH311). All synthesized cDNA was used as a template for PCR amplification of the region containing the *XBP1* splicing reporter and sgRNA sequence. The PCR products were prepared for paired-end sequencing on an Illumina platform (BerryGenomics). Raw FASTQ files for Read2 were cropped and aligned to the sgRNA reference of the library using Bowtie (v1.1.2) to identify sgRNAs, while Read1 files were cropped and aligned to a reference containing spliced and unspliced *XBP1* reporter sequences to determine splicing outcomes. The MAGeCK pipeline (v0.5.9.2)[68] was employed to determine sgRNA-level and gene-level phenotypes and significances.

### Immunoblotting

Cells were lysed in RIPA buffer (Beyotime, P0013B) supplemented with a phosphatase inhibitor cocktail (Selleck, B15001) and protease inhibitor cocktail (MCE, HY-K0010). The protein concentration of each sample was determined using the Easy II Protein Quantitative Kit (TransGen, DQ111). Proteins were denatured in loading buffer (Solarbio, P1041) by boiling at 95°C for 10 minutes, and 20 µg of protein from each lysate was electrophoresed on SDS-PAGE gels and transferred to 0.45 µm PVDF membranes (Immobilon, IPVH00010). The membrane was blocked with 5% BSA (Sigma-Aldrich, V900933) in TBST (Sangon, C520009) and incubated with primary antibodies as indicated at 4°C overnight. The next day, the membrane was incubated with horseradish peroxidase–conjugated secondary antibodies for 1 hour at room temperature. Protein signals were detected using the Clarity Western ECL Substrate (EpiZyme, SQ202L). The antibodies used in this study are summarized here: anti-PERK (Proteintech, 20582-1-AP), anti-Phospho-IRE1 (Abcam, ab124945), anti-IRE1 (CST, 3294), anti-GADD34 (Proteintech, 10449-1-AP), anti-XBP1 (CST, 12782), anti-ATF4 (Proteintech, 28657-1-AP), anti-Phospho-eIF2α (CST, 9721S), anti-eIF2α (CST, 5324), anti-GAPDH (Proteintech, HRP-60004), anti-Flag (Absin, Abs830014), anti-HA (Sigma, H3663), anti-GFP (Ray Antibody, RM1108).

### Co-immunoprecipitation (Co-IP)

3 × 10L HEK293T cells were plated onto a 10 cm diameter dish for 24 hours before transfection. The pcDNA3.1-GFP, pcDNA3.1-GFP-GADD34 FL, pcDNA3.1-GFP-GADD34 Δ1-60, pcDNA3.1-GFP-GADD34 Δ1-323, pcDNA3.1-GFP-GADD34 Δ513-674, PCDH-vector-3×Flag, and PCDH-FL GADD34-3×Flag vectors were transfected into the cells. Forty-eight hours later, the cells were collected and lysed in lysis buffer (Beyotime, P2181S-1) supplemented with Deoxy Big CHAP (BBI, A600681-0250). The cell lysates were centrifuged at 12,000 rpm at 4°C for 10 minutes, and the supernatant was collected. Anti-Flag magnetic beads (Beyotime, P2181S-4) and anti-GFP agarose beads (ABMagic, MA108-25T) were blocked overnight at 4°C using 5% BSA. The supernatant was then incubated with anti-Flag magnetic beads and anti-GFP agarose beads at 4°C overnight. After washing, the beads bound to proteins were denatured and subjected to immunoblotting analysis.

### Immunofluorescence (IF)

Cells were seeded at 5 × 10L cells per well on Matrigel-coated 12-mm glass coverslips (CITOTEST, #10210012CE) in 24-well plates and cultured overnight. Cells were then fixed with 4% paraformaldehyde (Beyotime, #P0099) for 30 min at room temperature (RT), permeabilized with 0.1% Triton X-100 (Coolaber, #CT11451) in PBS for 10 min, and blocked with 5% BSA (Sigma-Aldrich, #V900933) in PBS for 1 h at RT. Primary antibody against CALR (Proteintech, #10427-2-AP, 1:200 dilution) was applied overnight at 4°C. After washing three times with PBS (5 min each), cells were incubated with Alexa Fluor-conjugated secondary antibodies (1:500 dilution) for 1 hr at RT. Nuclei were counterstained with DAPI (Beyotime, #C1006, 1:1000) for 10 min at RT. Coverslips were washed three times with PBS and mounted using Mounting Medium (SouthernBiotech, #0100) on glass slides. Images were captured using a confocal microscope (Zeiss, LSM 980) and analyzed using Fiji (v2.0.0).

### Transduction of human T cells

Human CAR-T cells were generated as previously described[69]. Briefly, frozen peripheral blood mononuclear cells (PBMCs) from healthy donors were purchased from STEMCELL Technologies. Thawed PBMCs were activated by anti-CD3 (1 μg/mL, Biolegend, 317326) and anti-CD28 (1 μg/mL, BD Biosciences, 555725) mAbs pre-coated on a non-tissue culture plate. Activated T lymphocytes were then transduced with retroviral supernatants using retronectin-coated plates (Takara, T100B). Three days after transduction, CAR-T cells are harvested and cultured in complete human T cell media medium containing IL-7 (10 ng/mL; PeproTech, 200-07-500) and IL-15 (5 ng/mL; PeproTech, 200-15-500) for 7-9 days. 24hrs before functional assays, cytokines were washed away, and CAR-T cells were rested in T cell medium overnight.

### *Ex-vivo* exhaustion model

Tumor cells (BXPC3 or HCC1806-CD19) were seeded at 0.2 x 10^6^ cells per well in a 12 well plate overnight and B7-H3 or CD19 specific CAR-T cells were added in triplicate at 1-to-1 effector-to-target ratio in the presence of DMSO or 30 µM Sephin1 (MCE, HY-111022). 3 Days after co-culture, cells from all 3 wells were pooled, washed and resuspended in 3mL fresh media. 1mL of cells were used for cell counting and intracellular cytokine staining while 2mL of cells were added to freshly seeded tumor cells for additional rounds of co-culture.

### Intracellular cytokine staining

CAR-T cells were stimulated with 50 μg/mL PMA (Beyotime, S1819) and 1mM Ionomycin (Beyotime, S1672) with 5 μg/mL brefeldin A (Biolegend, 420601) for 4 hrs to induce cytokine production. Cells were then harvested and stained for CD8 and Live/Dead (Zombie Aqua, Biolegend, 423102) and subsequently fixed and permeabilized using BD Cytofix/Cytoperm™ Fixation/Permeabilization Kit (BD Bioscience, 554714) at 4 degrees overnight, followed by intracellular staining of IFN-γ and TNF-α.

### Flow cytometry analysis

The following flow cytometry antibodies were purchased from Biolegend: AF700-conjugated anti-CD8 (clone SK-1), AF647-conjugated anti-IFN-γ (clone 4S.B3), and PE-conjugated anti-TNF-α (clone MAB11). Flow cytometry data was acquired on a NovoCyte Quanteon flow cytometer (Agilent) using NovoExpress software and the flow data were analyzed by FlowJo software (v10.7, Tree Star).

## Acknowledgements

We thank the assistance of SUSTech Core Research Facilities on flow cytometry. This work was supported by the National Natural Science Foundation of China (82171416 to R.T., 32200769 to Y.X.), Shenzhen Medical Research Fund (A2303039 to R.T.), Guangdong Basic and Applied Basic Research Foundation (2023B1515020075 to R.T.) and Shenzhen Fundamental Research Program (JCYJ20220530112602006 and RCYX20221008092845052 to R.T., JCYJ20220530112604010 to Y.X.).

## Author contribution

R.T. conceived the project. R.T. and Y.X. supervised the project. Q.Y., Y.C. and L.S. conducted experiments and analyzed data with guidance from Y.X. and R.T. Q.Y., Y.C., Y.X. and R.T. wrote the manuscript with input from all authors.

## Supplementary files

Supplementary Figures 1-6 with legends

Supplementary Tables

— Supplementary Table 1: SPLiCR-seq screen results analyzed by the MAGeCK pipeline.

— Supplementary Table 2: Primer and sgRNA sequences used in this study.

