## Supplementary Figures for "SPLiCR-seq: A CRISPR-Based Screening Platform for RNA splicing Identifies Novel Regulators of IRE1-XBP1 Signaling Under ER Stress"

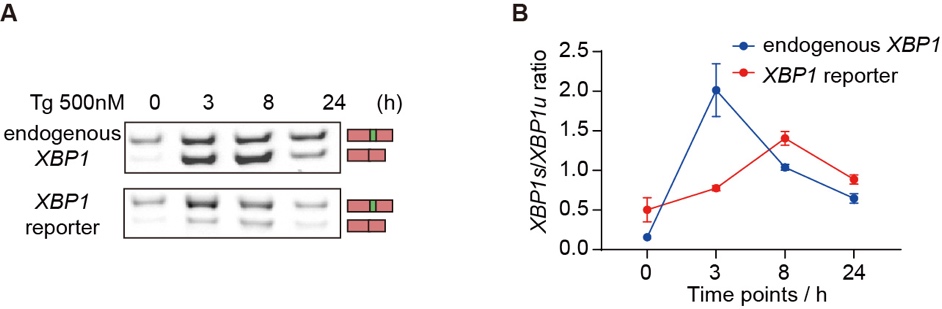

**Fig. S1: Dynamics of splicing of endogenous *XBP1* and XBP1 reporter in the SPLiCR-seq vector**

**(A)**	Time-course analysis of *XBP1* splicing dynamics for endogenous *XBP1* and the XBP1 reporter in the SPLiCR-seq vector in HEK293T cells treated with Tg (500 nM) for the indicated durations, as assessed by RT-PCR. Spliced and unspliced products are indicated.

**(B)** Quantification of time-course splicing levels for endogenous *XBP1* (blue, mean ± SEM, n = 6 biological replicates) and XBP1 reporter (red, mean ± SEM, n = 2 biological replicates) using grayscale analysis of RT-PCR bands.

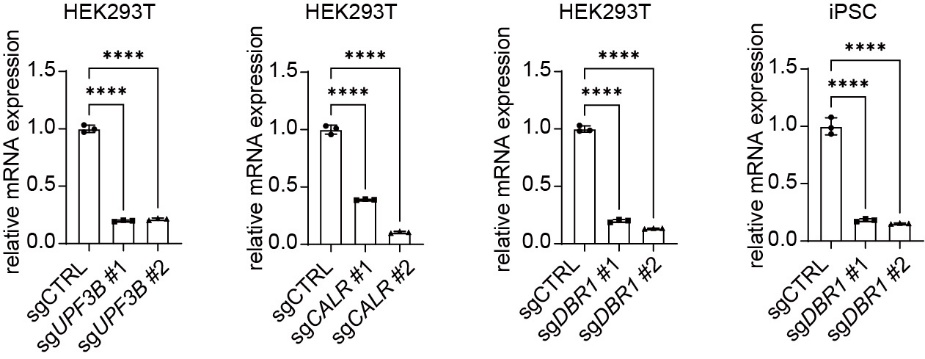

**Fig. S2: Validation of Gene Knockdown in CRISPRi-HEK293T Cells and CRISPRi-iPSCs**

qPCR quantification of mRNA expression levels for *UPF3B*, *CALR*, and *DBR1* in CRISPRi-HEK293T cells and CRISPRi-iPSCs transduced with sgRNAs targeting each gene. A non-targeting sgRNA (sgCTRL) was used as the reference. Data are shown as mean ± SD (n = 3 technical replicates). Statistical analysis was performed using one-way ANOVA (****P < 0.0001).

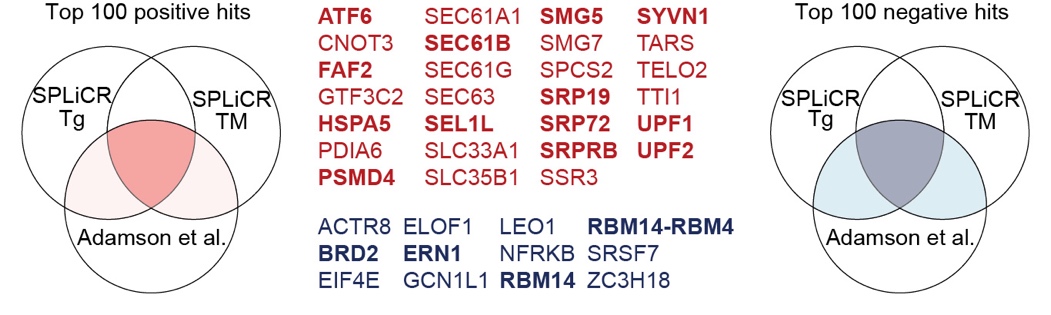

**Figure S3: Overlap Analysis of SPLiCR-seq and a Previous XBP1 Screen**

Venn diagrams comparing the top 100 positive (left) and negative (right) hits identified in our genome-wide SPLiCR-seq screens under Tg and Tm treatments with those reported in Adamson et al. Common genes identified between the Adamson et al. screen and either SPLiCR screen are shown in red (positive hits) or blue (negative hits). Genes shared across the Adamson et al. screen and both SPLiCR screens are highlighted in bold.

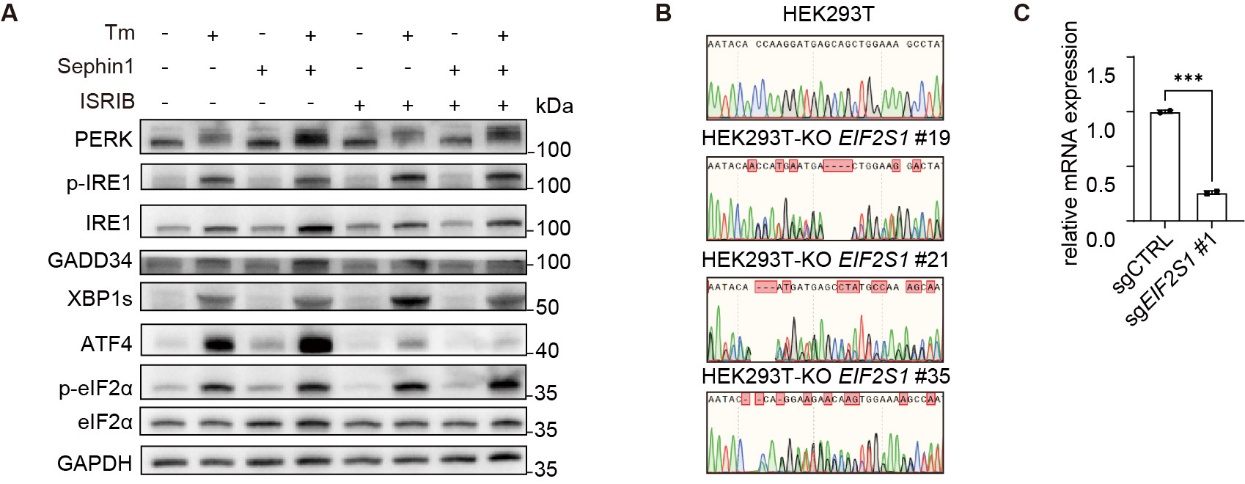

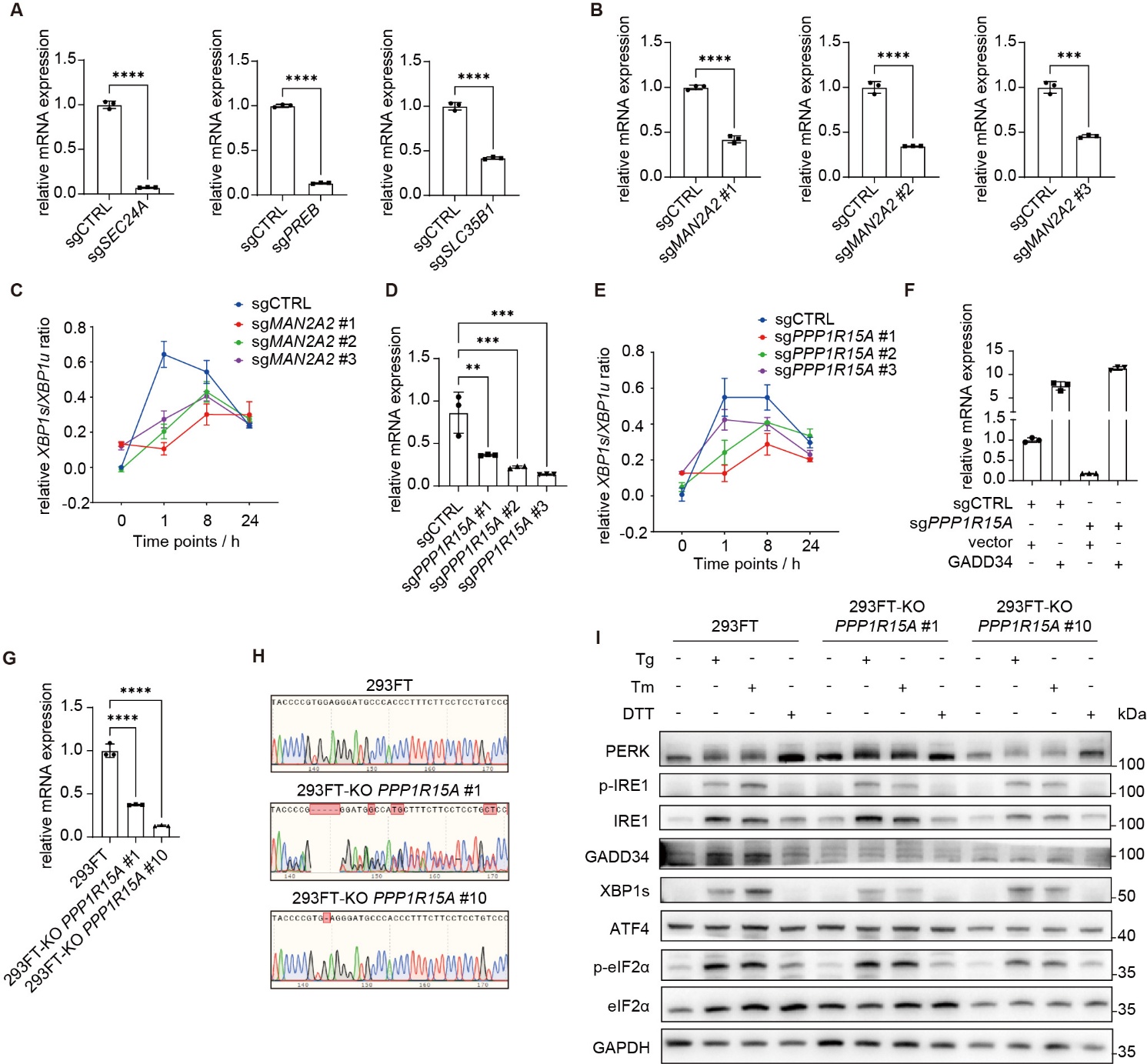

**Fig. S4: Validation of screening hits**

**(A)** qPCR quantification of mRNA expression levels for *SEC24A*, *PREB*, and *SLC35B1* in CRISPRi-HEK293T cells transduced with sgRNAs targeting each gene. A non-targeting control sgRNA (sgCTRL) was used as the reference. Data are shown as mean ± SD (n = 3 technical replicates). Statistical analysis was performed using Student’s t-test (****P < 0.0001).

**(B)** qPCR quantification of mRNA expression levels for *MAN2A2* in CRISPRi-HEK293T cells transduced with three different sgRNAs targeting *MAN2A2*. A non-targeting control sgRNA (sgCTRL) was used as the reference. Data are shown as mean ± SD (n = 3 technical replicates). Statistical analysis was performed using Student’s t-test (***P<0.001, ****P<0.0001).

**(C)** Time-course qPCR analysis of *XBP1* splicing levels in control and *MAN2A2* knockdown CRISPRi-HEK293T cells treated with Tg (500 nM) for the indicated durations. Data are shown as mean ± SD (n = 3 technical replicates).

**(D)** qPCR quantification of mRNA expression levels for *PPP1R15A* in CRISPRi-HEK293T cells transduced with three different sgRNAs targeting *PPP1R15A*. A non-targeting control sgRNA (sgCTRL) was used as the reference. Data are shown as mean ± SD (n = 3 technical replicates). Statistical analysis was performed using one-way ANOVA (**P<0.01, ***P<0.001, ****P<0.0001).

**(E)** Time-course qPCR analysis of *XBP1* splicing levels in control and *PPP1R15A* knockdown CRISPRi-HEK293T cells treated with Tg (500 nM) for the indicated durations. Data are shown as mean ± mean ± SD (n = 3 technical replicates).

**(F)** qPCR quantification of *PPP1R15A* (GADD34) expression in control (sgCTRL) and *PPP1R15A*-knockdown (sgPPP1R15A) CRISPRi-HEK293T cells overexpressing either an empty vector or a GADD34-Flag vector. Data are shown as mean ± standard deviation (n = 3 technical replicates). sgCTRL cells transduced with the empty vector were used as the reference.

**(G)** qPCR quantification of mRNA expression levels for *PPP1R15A* in *PPP1R15A* knockout 293FT cells. Wild type 293FT was used as the reference. Data are shown as mean ± SD (n = 3 technical replicates). Statistical analysis was performed using one-way ANOVA (****P<0.0001).

**(H)** Sanger sequencing electropherograms showing the editing of *PPP1R15A* in *PPP1R15A* knockout 293FT cells, confirming the knockout of *PPP1R15A*.

**(I)** Western blot analysis of PERK, p-IRE1, total IRE1, GADD34, XBP1s, p-eIF2α, total eIF2α and ATF4 in WT and *PPP1R15A* knockout 293FT cells treated with Tg (500 nM, 24 h), Tm (2.5 μM, 24 h), or DTT (2 mM, 24 h). GAPDH was used as a loading control.

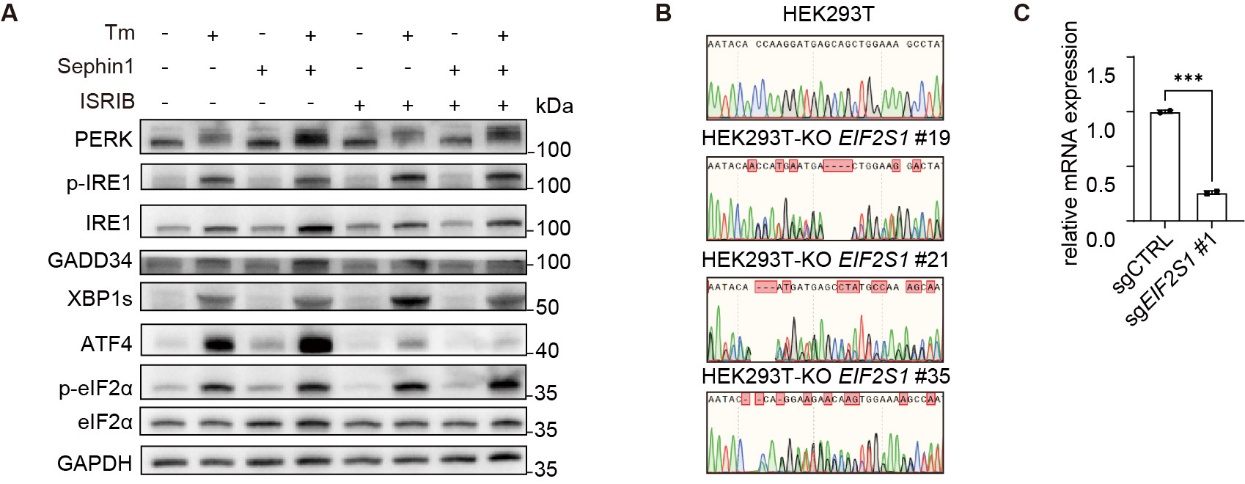

**Fig. S5: Sephin1 Inhibits the IRE1-XBP1 Pathway Independently of eIF2α**

**(A)** Western blot analysis of PERK, p-IRE1, total IRE1, GADD34, XBP1s, p-eIF2α, total eIF2α and ATF4 in HEK293T cells treated with Tm (2 μM, 24 h), Sephin1 (5 μM, 24 h), or ISRIB (1 μM, added 3 h before cell collection) as indicated. GAPDH was used as a loading control.

**(B)** Sanger sequencing electropherograms showing the editing of *EIF2S1* in *EIF2S1* knockout 293FT clones, confirming the partial knockout of *EIF2S1*.

**(C)** qPCR quantification of *EIF2S1* mRNA expression in wild-type control (sgCTRL) and *EIF2S1* knockout 293FT cells. Data are represented as mean ± SD (n = 3 technical replicates). Wild-type control was used as the reference. Statistical analysis was performed using Student’s T-test (***P<0.001).

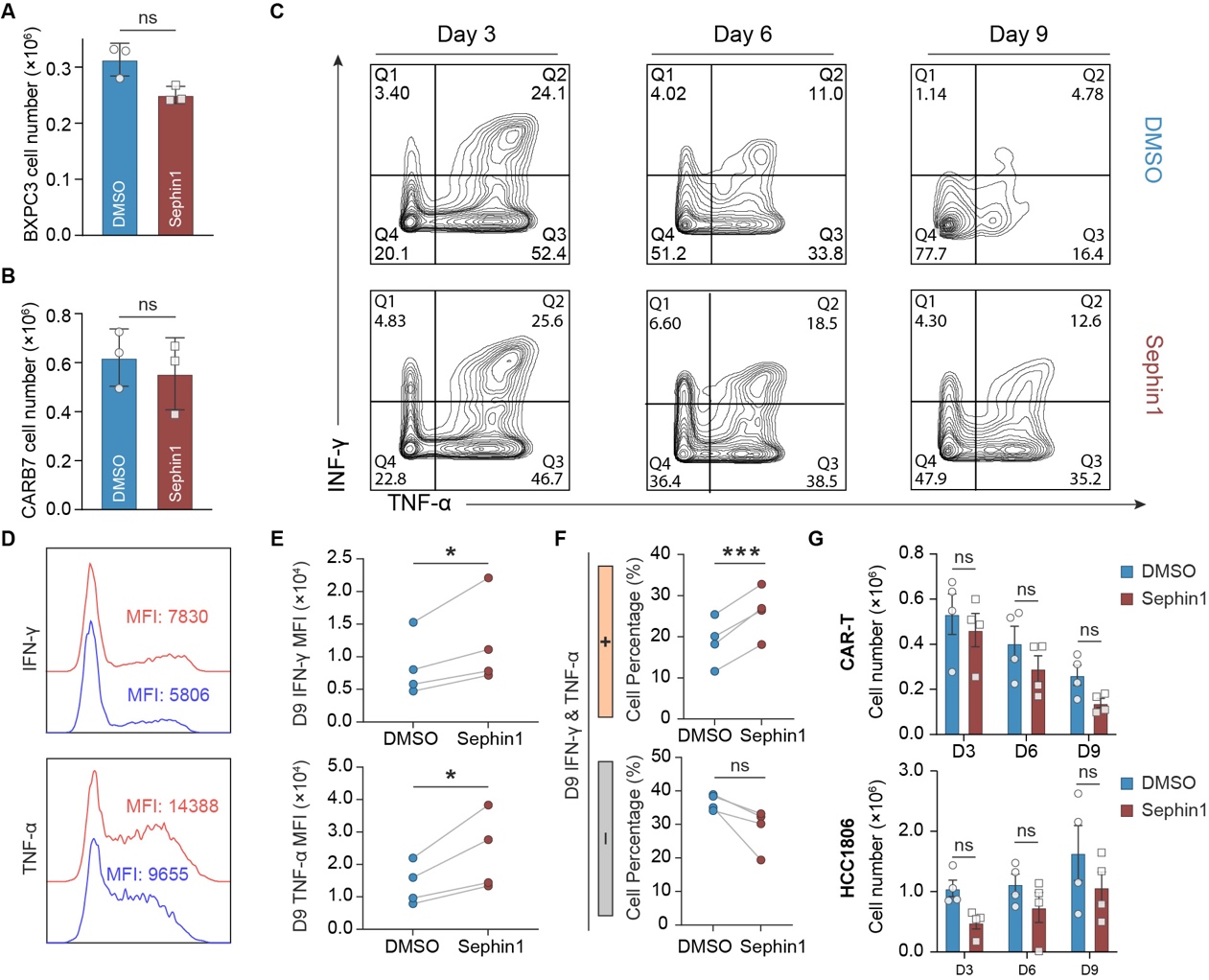

**Fig. S6: Sephin1 enhances cytokine secretion capacity of CD19 CAR-T cells in *ex-vivo* exhaustion model**

**(A, B)** Cell count of BXPC3 cells (**A**) and CAR-T Cells (**B**) treated with DMSO or Sephin1 (Data are presented as mean ± SEM, paired Student’s t-test; *P < 0.05, n=3 experiments).

**(C, D)** Intracellular cytokine staining of IFN-γ and TNF-α in CAR-T cells repetitively co-cultured with tumor cells for 3, 6 and 9 days in the presence of DMSO or Sephin1 and the percentage of IFN-γ^+^/TNF-α^+^ and IFN-γ^-^/TNF-α^-^ in CAR-T cells at day 9.

**(E, F)** Expression of of IFN-γ and TNF-α in CAR-T cells at day 9 and the mean fluorescent intensity (MFI) of IFN-γ and TNF-α in cells stained positive for respective cytokine. (Data are presented as mean ± SEM, paired Student’s t-test; *P < 0.05, n=4 experiments).

**(G)** Cell counts of CAR-T cells and HCC1806-CD19 cells at day 3, 6, 9 upon treatment of DMSO or Sephin1 (Data are presented as mean ± SEM, two-way ANOVA with Sidak multiple comparisons test; ns, not significant, *P < 0.05, **P < 0.01, n=4 experiments).
